# Leaf functional trait evolution and its putative climatic drivers in African *Coffea* species

**DOI:** 10.1101/2024.02.13.580072

**Authors:** Aiden Hendrickx, Yves Hatangi, Olivier Honnay, Steven B. Janssens, Piet Stoffelen, Filip Vandelook, Jonas Depecker

## Abstract

**Background and Aims:** Leaf traits are known to be strong predictors of plant performance and can be expected to (co)vary along environmental gradients. We investigated the variation, integration, environmental relationships, and evolutionary history of leaf functional traits in the genus *Coffea* L., typically a rainforest understory shrub, across Africa. A better understanding of the adaptive processes involved in leaf trait evolution can inform the use and conservation of coffee genetic resources in a changing climate.

**Methods:** We used phylogenetic comparative methods to investigate the evolution of six leaf traits measured from herbarium specimens of 58 African *Coffea* species. We added environmental data and data on maximum plant height for each species to test trait-environment correlations in various (sub)clades, and we compared continuous trait evolution models to identify variables driving trait diversification.

**Key Results:** A substantial leaf trait variation was detected across the genus *Coffea* in Africa, which was mostly interspecific. Of these traits, stomatal size and stomatal density exhibited a clear trade-off. We observed low densities of large stomata in early branching lineages and higher densities of smaller stomata in more recent taxa, which we hypothesise to be related to declining CO_2_ levels since the mid-Miocene. Brownian Motion evolution was rejected in favour of White Noise or Ornstein-Uhlenbeck models for all traits, implying these traits are adaptively significant rather than driven by pure drift. The evolution of leaf area was likely driven by precipitation, with smaller leaves in dryer climates across the genus.

**Conclusions:** Generally, *Coffea* leaf traits appear to be evolutionarily labile and governed by stabilising selection, though evolutionary patterns and correlations differ depending on the traits and clades considered. Our study highlights the importance of a phylogenetic perspective when studying trait relationships across related taxa, as well as the consideration of various taxonomic ranges.

## Introduction

Since the emergence of the first vascular plants over 400 mya, evolution has seen the unfolding of a remarkable diversity of leaf traits (Hao & Xue, 2013). Morphologically, leaves are composed of various traits presenting different degrees of appearance. Since specific leaf trait combinations are known to have evolved in synchrony with a plant’s survival strategy, leaf traits strongly predict a plant’s performance in the environment it grows in (Poorter & Bongers, 2006; Violle et al., 2007). Since traits are often phenotypically integrated, several leaf functional traits have been compiled into the Leaf Economic Spectrum (LES) (Wright et al., 2004). This framework represents a continuum of survival strategies, ranging from fast to slow return on investment in leaf biomass (Wright et al., 2004; Jones et al., 2013; Reich, 2014). Furthermore, because functional traits form the interface between the plant and its surroundings, evolutionary change in these traits facilitates the adaptation of plants to a changing environment (Ackerly et al., 2000; Ackerly, 2004; Jones et al., 2013).

Leaf functional traits, by definition, have the capacity to influence the performance or fitness of an organism in its environment (Reich et al., 2003; McGill et al., 2006). They have been related to performance measures such as Water Use Efficiency (WUE) (e.g., Xu & Zhou, 2008; Drake et al., 2013) and tend to covary with environmental variables, such as temperature, precipitation, and nutrient availability (Givnish, 1987; Wright et al., 2017). Leaf size, for instance, is often larger in hot and wet tropical climates, while smaller leaves predominate in cooler or more arid regions (Wright et al., 2017). Plants have also shown a reduced stomatal size and increased stomatal density when exposed to drought stress (Xu & Zhou, 2008), though this response is likely species-specific (Zhang et al., 2012). Further, specific leaf area (SLA, i.e., leaf area per unit of dry mass) has been found to increase with increasing soil nutrient levels (Andersen et al., 2012) and soil moisture content (Chaturvedi & Raghubanshi, 2018) in tropical forest species.

Despite the wealth of research focusing on leaf trait evolution at different taxonomic levels (e.g., Flores et al., 2014; Glade-Vargas et al., 2018), sources of trait variation often remain poorly documented, particularly in tropical forest understories. The lack of data leads to gaps in our understanding of the adaptive potential of tropical taxa. A closer look at particularly understudied floras, such as the Afrotropical understory, is required to gain insights into the current state and evolutionary history of these ecosystems (Verbeeck et al., 2011). Broad studies of leaf trait evolution across Africa are lacking, though some studies have been performed at smaller geographical scales (e.g., Lauterbach et al., 2016; Wigley et al., 2016). As the understory of African tropical forests harbours important crop wild relatives, including numerous wild coffee species, an enhanced focus on these species is also crucial for the long-term conservation of their genetic resources. Although some research has been done on leaf trait diversity in the Congo Basin canopy layer (Verbeeck et al., 2014; Kafuti et al., 2020), knowledge on leaf traits of understory species remains very limited (but see Hatangi et al., 2023).

Coffee is a valuable commodity worldwide, with global consumption exceeding ten million tons annually (International Coffee Organization, 2023). The popular beverage is brewed from seeds of a few *Coffea* species; *C. arabica* L. and *C. canephora* Pierre ex A.Froehner make up the vast majority of the global coffee market, which exceeds a total value of 88 billion US dollars (International Coffee Organization, 2023; Statista, 2023). Despite this duopoly in the global market, other *Coffea* species, such as *C. liberica* and *C. stenophylla*, have shown promise as alternative beverage species (Davis et al., 2021, 2022). The genus *Coffea* encompasses 131 species known to date, after the inclusion of 20 species formerly classified as a sister genus, *Psilanthus* (Robbrecht & Manen, 2006; Davis et al., 2007, 2011; Hamon et al., 2017) and the recent description of seven new species (Davis & Rakotonasolo, 2021; Stoffelen et al., 2021). Most *Coffea* species are native to continental Africa or Madagascar, with a few species occurring in Comoros and Mayotte, the Mascarene islands, in parts of Asia, or in Northern Australia (Davis, 2003; Hamon et al., 2017). Most of these species have relatively narrow geographic ranges (Davis et al., 2006). Nonetheless, some species such as *C. canephora* and *C. liberica* occur across large areas of the African continent (Herrera & Lambot, 2017). These widespread species tend to inhabit moist, tropical habitats, or areas along rivers or wetlands. In contrast, other *Coffea* species can be found in dryer shrublands, or in deciduous forests with a distinct dry season (Maurin et al., 2007; Herrera & Lambot, 2017).

Two major clades have been identified within the genus: a relatively small xeno-coffee (XC) clade, and a more diverse eu-coffee (EC) clade (Hamon et al., 2017). The sparsity of the XC clade is assumed to be related to haploidy and selfing within the clade, whereas nearly all EC species are diploid and self-incompatible (Davis et al., 2005; Hamon et al., 2017). The XC clade is dispersed over tropical West, Central and East-Africa as well as Asia and Northern Australia, while the EC clade encompasses two subclades occurring across continental tropical Africa (AFR subclade) and the West Indian Ocean Islands (WIOI subclade) (Davis, 2003; Hamon et al., 2017). Extant taxonomic diversity has been influenced by several environmental and geographic factors. For example, the Dahomey Gap, an arid savannah region separating West and Central African forests, is known to have influenced the genetic diversity and speciation within *Coffea* by acting as a barrier to gene flow (Berthaud, 1986; Maurin et al., 2007; Gomez et al., 2009; Cubry et al., 2013). Further, speciation on the West Indian Ocean Islands has been shown to be rapid and radial, suggesting a possible adaptive radiation in the WIOI subclade (Davis et al., 2006; Anthony et al., 2010). Learning more about the macroevolution of leaf traits in these species, specifically in response to for example climatic variables, can inform their conservation to safeguard the genetic resources within these species.

Previous studies investigating leaf trait variation in *Coffea* have focused on only one or a few species (e.g., Buchanan et al., 2019; Dutra Giles et al., 2019; Dubberstein et al., 2021). Moreover, although several studies have assessed the macroevolution of leaf functional traits across a given taxonomic group (e.g., Flores et al., 2014; Onstein et al., 2016; Glade-Vargas et al., 2018), the drivers and mode of leaf trait evolution across a principally rainforest understory genus such as *Coffea* remains unexplored. For example, a different pattern of leaf trait evolution can be expected between continental African (AFR) and Madagascan (WIOI) species, due to their independent evolutionary histories and different biogeographical configuration (Hamon et al., 2017). Furthermore, species that have adapted to survive in drier habitats could be expected to reflect these adaptations in their leaf traits, for example via lower SLA values (Chaturvedi & Raghubanshi, 2018). Also, functional trait variation is a prerequisite for natural selection and adaptive trait change. We expect interspecific trait variation to exceed intraspecific variation for all traits, due to most *Coffea* species’ fairly narrow distribution range (Davis et al., 2006). We also expect some traits to show a certain degree of integration. For example, stomatal density and stomatal size are generally inversely correlated across taxa (Brodribb et al., 2013). Overall, we predict that trait-trait correlations will be stronger than trait-environment relationships, due to the large amount of trait variation found between coexisting species (Wright et al., 2004; Moles et al., 2005; Vandelook et al., 2012; Jones et al., 2013). Finally, by evaluating whether trait evolution occurred independently from the phylogenetic relationships between taxa, it is possible to shed light on the importance of adaption and constraints in trait evolution. At present, these hypotheses have remained untested in *Coffea*. Current knowledge on evolutionary drivers in *Coffea* is incomplete, and a better understanding of the processes involved in trait evolution could shed light on the adaptive value of the studied leaf traits in wild coffee plants. Additionally, knowledge on trait-trait and trait-climate relationships is essential to foster a more resilient and sustainable coffee cultivation.

Via the use of phylogenetic comparative methods, we aim to (i) estimate the relative degree of intra- and interspecific leaf functional trait variation across 58 African *Coffea* species that are broadly distributed across the genus, across Africa, and across habitats; (ii) analyse potential correlated evolution between traits; and (iii) explore which environmental factors are the most likely drivers of leaf trait evolution across the genus. We also attempt to discern different life history strategies among *Coffea* clades or species and to identify the evolutionary trajectories that led to the current trait diversity in the genus.

## Materials & methods

### Data collection

A data set of trait measurements was compiled from 780 leaves across 167 herbarium accessions of 62 species. Most of the included species (n = 58) belonged to the genus *Coffea*, except for four outgroup species from related genera (*Belonophora coriacea*, *Calycosiphonia spathicalyx*, *Tricalysia congesta,* and *Bertiera iturensis*). Of the 58 *Coffea* species, two were representatives of the former *Psilanthus* genus. The samples included in this study were well distributed throughout the genus, across habitats, and across Africa. The leaf functional traits included in the analyses were leaf area (m^2^), SLA (m^2^ kg^-1^), stomatal density (mm^-2^), stomatal length (µm), stomatal width (µm), and pore width (i.e., aperture width, µm). Data on leaf dry mass and pore length was also compiled, but these traits were not further considered due to strong correlations with leaf area (*r_s_* = 0.898, P < 0.001) and stomatal length (*r_s_* = 0.817, P < 0.001), respectively. Leaf area was measured by scanning leaves using a standard A4 flatbed scanner and estimating area using ImageJ (Schneider et al., 2012). Leaf dry mass was determined by weighing individual leaves, after a week of calibration to laboratory moisture conditions (50 to 60% RH). To correct for potential bias in SLA estimates due to shrinkage during the drying of herbarium specimens (Blonder et al., 2012; Perez et al., 2020), we calculated average shrinkage factors for species of which live specimens were available in Meise Botanic Garden greenhouses. The methodology of this correction is presented in the Appendix.

For the same leaves, stomatal prints were taken from the abaxial side using the nail varnish method as explained in Meeus et al. (2020). Two prints per leaf were taken in the middle of the leaf at opposite sides of the main vein, from which three photomicrographs of 1,600 × 1,200 pixels were taken per leaf print (dimensions = 344 × 258 µm; area view field = 0.09 mm^2^) using a digital microscope (VH-5000 Ver 1.5.1.1, KEYENCE CORPORATION). A single photomicrograph was created by stacking several digital images taken at different focal planes to increase the depth of the resulting image. All stomata that fell entirely within the view field were counted and converted to stomata per square millimetre to obtain stomatal density. Stomatal measurements were performed on the same digital photos using ImageJ version 1.53v (Schneider et al., 2012) on three stomata per leaf across five leaves (i.e. 15 stomata) per accession. Stomatal dimensions were not measured for three species (*C. homollei*, *C. arenesiana*, and *C. coursiana*) due to a lack of source material, and one leaf was removed from the data set due to an exceptionally low leaf weight at around five times the area of the other leaves from this accession (specimen Vermoesen 2182, **[Supplementary information]** Table S1). This yielded stomatal measurements of 742 leaves representing 159 accessions across 59 species (55 *Coffea* + 4 outgroup species). Trait values for all leaves were averaged per species to obtain a comprehensive data set including a single mean trait value per species (**[Supplementary information]** Table S2). Standard errors were added to the averaged trait data, and leaf area was log-transformed before analysing the data to avoid violating normality of residuals.

Additionally, data on maximum plant height for each species was compiled from various literature sources (Leroy, 1961; Bridson, 1994; Stoffelen, 1998; Davis & Rakotonasolo, 2001, 2008; Ruffo et al., 2002; Davis & Mvungi, 2004; Davis et al., 2006). These values were added to the data set, along with geographic coordinates and altitude of all accessions. Geographic coordinates were obtained by georeferencing based on the locality information provided on the specimens, which was detailed up to at least the nearby village level.

Climate data for each accession was retrieved from WorldClim (Fick & Hijmans, 2017) for each of the given coordinates in the form of 19 bioclimatic variables, which were subsequently averaged at species level. To reduce the dimensionality of these variables, a phylogenetically corrected PCA was performed on the correlation matrix to decompose the climate data into phylogenetically structured principal components (PCs) using R package *phytools* (Revell, 2012). Three interpretable PCs were retained, jointly explaining 79% of the variance in the climate data. These three PCs were added to the dataset and used later as predictors in the phylogenetic regression analyses and evolutionary models.

### Phylogenetic tree construction

(R Core Team, 2023)Phylogenetic information included in the analyses was based on the phylogeny of Hamon et al. (2017). We pruned their phylogeny to retain only those species for which data were readily available to us, using the *drop.tip* function in R package *ape* (Paradis & Schliep, 2019). The full tree including outgroups was ensured to be ultrametric (using the *chronos* command in *ape* with a smoothing parameter of zero) and dichotomous (using *multi2di* in *ape*) before rescaling its length to 1. Subtrees were then extracted from the full phylogeny to include (i) only *Coffea* species, (ii) only the EC clade, (iii) only the AFR subclade, and (iv) only the WIOI subclade. The XC clade was not extracted to a separate subtree due to its small size. These subtrees allowed for the isolation of different clades, to then run independent regression analyses for each of them (see section “Phylogenetic regressions”). All analyses were performed in R version 4.3.0 (R Core Team, 2023).

### Inter- and intraspecific variation

The proportion of inter- and intraspecific variation in the trait values was estimated using one-way Analysis of Variance (ANOVA). Although the assumption of independence was violated, the results are presented heuristically. We then applied a phylogenetically corrected PCA to the trait data to identify axes of multivariate trait evolution using *phytools* (Revell, 2012). The results were visualised in biplots, as well as in phylomorphospace plots as implemented in *phytools* to visualise phylogenetic patterns.

### Phylogenetic signal

We estimated Pagel’s lambda (λ) (Pagel, 1999), a measure of phylogenetic signal in the trait data, with the *phylosig* command in *phytools* while accounting for standard error in the data. Pagel’s λ varies between 0 and 1, with 0 indicating phylogenetically independent trait evolution, and 1 indicating the trait followed a perfect Brownian model of evolution. Signal was estimated in all considered clades and subclades.

### Models of continuous trait evolution

To test whether trait evolution had been driven by environmental variables, we fitted different models reflecting different evolutionary processes. These models allow interpretations of the evolutionary processes involved in the diversification of *Coffea* leaf functional traits. The first and simplest model used was a non-phylogenetic “white noise” (WN) model, which assumes phylogenetic independence (λ = 0). This model is equivalent to drawing traits randomly from a normal distribution, independently of the phylogeny (Pagel, 1999; Münkemüller et al., 2015). The second model was simple Brownian Motion (BM), which is a constant-variance random-walk model that assumes that λ = 1 and that traits change randomly over time at a given rate (Freckleton et al., 2002; Revell et al., 2008; Meireles et al., 2020). This model implies that the evolution of the trait has been driven by pure drift, though the same pattern could be found under natural selection fluctuating in direction and intensity through time (O’Meara et al., 2006; Losos, 2008). Third, we fitted an Ornstein-Uhlenbeck (OU) model (Hansen, 1997; Butler & King, 2004; Hansen et al., 2008), which can be used to model evolution toward an optimal trait value *θ*. OU models contain a stochasticity parameter, *σ^2^*, which expresses the intensity of random fluctuations in trait values, and an adaptive parameter *α*, which measures the rate of trait change toward the optimum. An OU model with a single optimum can represent constrained evolution or inertia. WN models can also be described as variations on OU models, but with *α* = ∞ (Münkemüller et al., 2015). If leaf trait evolution is adaptive, a White Noise (WN) or Ornstein-Uhlenbeck (OU) model of evolution should be the best fit to the data (Hansen, 1997; Butler & King, 2004; Beaulieu et al., 2012; Münkemüller et al., 2015).

Since optima are expected to vary with evolutionary drivers such as climate, we also fitted four OU models in which the evolutionary optimum varies continuously as a function of a given putative driver of trait evolution (OU_A_ = altitude; OU_T_ = temperature, i.e., climate PC1; OU_D_ = drought, i.e., climate PC2; OU_S_ = Seasonality, i.e., climate PC3). In all OU models, we also estimated phylogenetic half-life (*t_1/2_* = ln(2)/*α*), another measure of phylogenetic signal (Hansen, 1997). This metric represents the time required for the average trait value to move halfway towards the optimum *θ*.

The WN models were fitted using the *fitContinuous* function in R package *geiger* (Pennell et al., 2014). We used the *brown.fit* and *slouch.fit* commands in R-package *slouch* (Kopperud et al., 2020) to fit the BM and OU1 models, as well as the more complex OU_T_, OU_D_, OU_S_, and OU_A_ models. Model fit was assessed using AICc, and measurement error was always incorporated into the models.

### Phylogenetic regressions

To test for correlated evolution between traits and for any effects of environmental variables on trait evolution, we fitted regression models for each leaf trait across four different phylogenies (i.e., the genus *Coffea*, the EC clade, the AFR subclade, and the WIOI subclade), each time with all other traits and environmental variables (i.e., Climate PC1, PC2, PC3, altitude, latitude, and longitude) as predictors. Since leaf traits are known to vary with plant size in some taxa (Price et al., 2014), maximum plant height was also included as a predictor in the regression models to account for possible allometric relationships between leaf traits and plant size.

To account for any non-independence in the species data due to common ancestry, we used phylogenetic generalised least squares (PGLS) regressions with the *pgls* command from R package *caper* (Orme et al., 2018). Applying a phylogenetic correction to a linear regression based only on univariate estimates of phylogenetic signal can lead to erroneous inferences (Hansen & Orzack, 2005; Revell, 2010). Therefore, *pgls* estimates phylogenetic signal simultaneously with the regression parameters. A backward model selection approach was used to remove the least significant predictors one by one until model AICc was minimal.

## Results

### Distribution of variance

Interspecific trait differences were significant for all traits and explained most of the variance for all traits except pore width (Table 1, **[Supplementary information]** Figure S1). The phylogenetic PCA of the trait data resulted in two meaningful PC axes, jointly explaining 64% of the variance in the trait data (Figure 1A, B). Trait PC1 showed strong negative loadings for stomatal dimensions (stomatal length: -0.94, stomatal width: -0.86, pore width: -0.70) and a substantial positive loading for stomatal density (-0.57) **[Supplementary information]** Table S3). PC axis 1 can therefore be regarded as a continuum from species with large stomata (generally at lower densities) to species with small stomata (generally at higher densities), explaining 42% of the trait variance. The second trait PC axis had a negative loading for SLA (-0.66) and positive loadings for stomatal density and leaf area (both +0.63), explaining another 22% of the trait variance. This PC axis therefore represents a spectrum from low leaf investment (small leaves, lower stomatal density, high SLA) to high leaf investment (larger leaves with higher stomatal density and low SLA).

**Figure 1:**
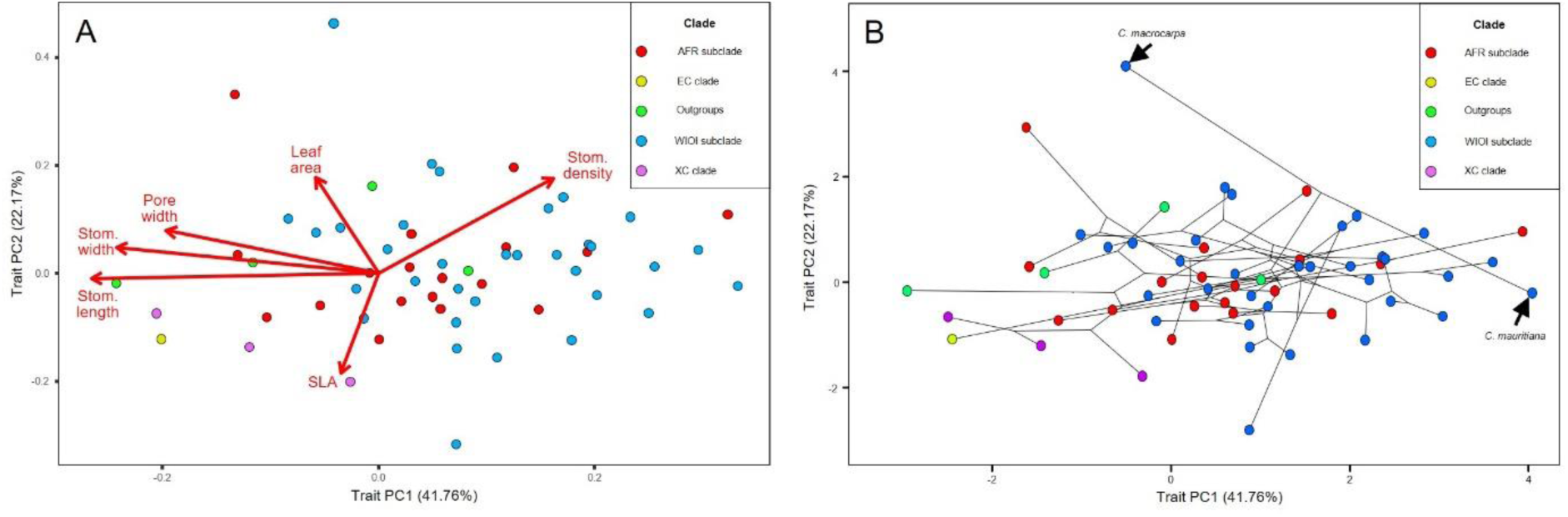
Principal component analysis showing the relationships between leaf traits, showing the first two PC axes. Data points indicate species (n = 62) and are color-coded according to their clade. **A**, biplot; **B**, phylomorphospace plot with lines representing phylogenetic relationships. AFR = African; EC = Eu-coffee; WIOI = West Indian Ocean Islands; XC = Xeno-coffee. Clade nomenclature is based on Hamon et al. (2017).

**Table 1:**
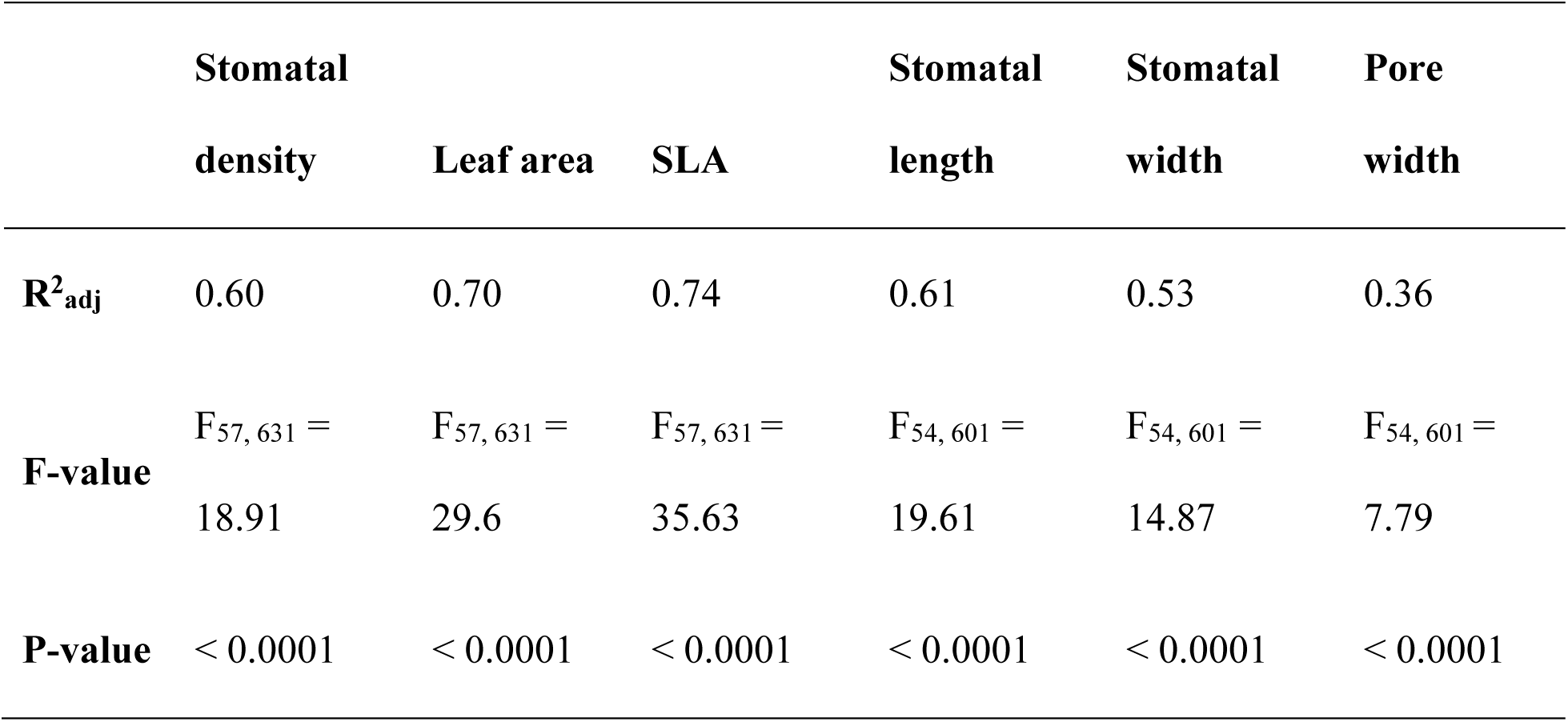
Variance explained by between-species differences for each studied leaf trait across *Coffea*, estimated from one-way Analysis of Variance (ANOVA). For stomatal density, leaf area, and SLA, n = 58; for stomatal length, stomata width, and pore width, n = 55. R^2^_adj_ = Adjusted R^2^.

|  | **Stomatal**<br>**density** | **Leaf area** | **SLA** | **Stomatal**<br>**length** | **Stomatal**<br>**width** | **Pore**<br>**width** |
| --- | --- | --- | --- | --- | --- | --- |
| **$R^2_{\text{adj}}$** | 0.60 | 0.70 | 0.74 | 0.61 | 0.53 | 0.36 |
| **F-value** | $F_{57, 631} =$<br>18.91 | $F_{57, 631} =$<br>29.6 | $F_{57, 631} =$<br>35.63 | $F_{54, 601} =$<br>19.61 | $F_{54, 601} =$<br>14.87 | $F_{54, 601} =$<br>7.79 |
| **P-value** | < 0.0001 | < 0.0001 | < 0.0001 | < 0.0001 | < 0.0001 | < 0.0001 |

### Univariate phylogenetic signal

Within the subclades of the genus *Coffea*, none of the tested traits showed significant phylogenetic signal except stomatal density in the WIOI subclade (λ = 0.69, P = 0.015) (Figure 2). Phylogenetic signal in leaf area was marginally significant across the EC clade (λ = 0.55, P = 0.087). Across the genus, significant signal was detected for SLA (λ = 0.52, P = 0.008) and stomatal length (λ = 0.73, P = 0.006), while leaf area remained marginally significant (λ = 0.69, P = 0.075). When including outgroups, phylogenetic signal in stomatal width became significant (λ = 0.57, P = 0.006), along with SLA (λ = 0.54, P = 0.018) and stomatal length (λ = 0.75, P = 0.001). All other traits were randomly distributed across the considered clade, with more related species not being significantly more similar to each other than less related lineages.

**Figure 2:**
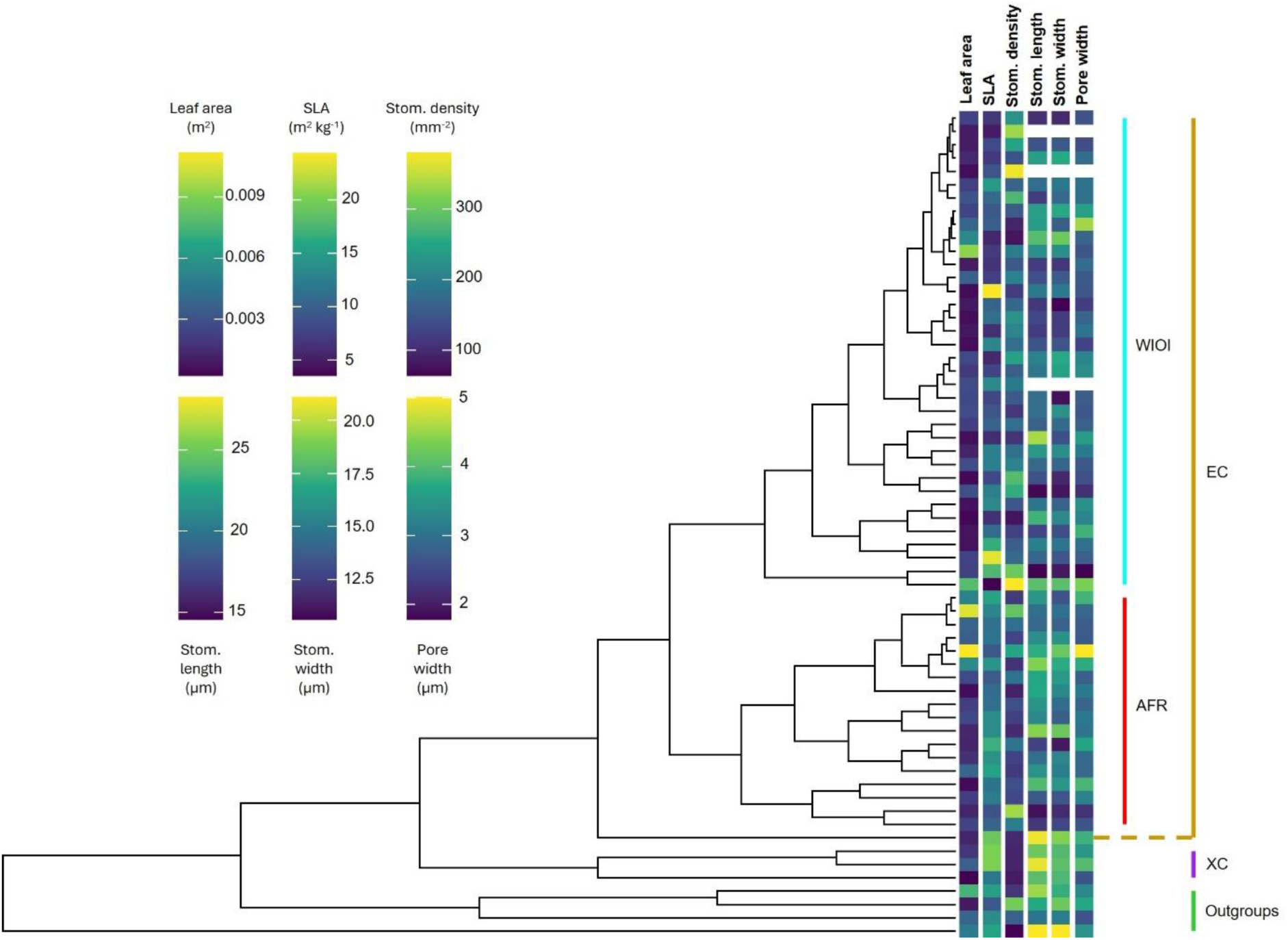
Phylogenetic tree of all studied species (n = 62) with values for the six considered traits shown at the tips of the tree. For stomatal length, stomatal width and pore width, trait values were missing for *C. arenesiana*, *C. coursiana*, and *C. homollei*. Clades are delimited by the bars at the right-hand side. WIOI = West Indian Ocean Islands; AFR = African; EC = Eu-coffee; XC = Xeno-coffee. Clade nomenclature is based on Hamon et al. (2017).

### Multivariate phylogenetic signal

The PGLS regressions did not detect phylogenetic signal (λ) significantly different from zero in any of the considered leaf traits across *Coffea* or across the EC clade, despite yielding non-zero maximum likelihood (ML) estimates for some traits (Table 2, **[Supplementary information]** Table S4). However, ML estimates for λ became significantly different from 0 for some traits across the subclades. For stomatal density and stomatal length in the AFR subclade, λ = 1. In the WIOI subclade, signal for stomatal density was strong, but significantly below 1 (λ = 0.945; 95% CI = 0.801, 0.992).

**Table 2:**
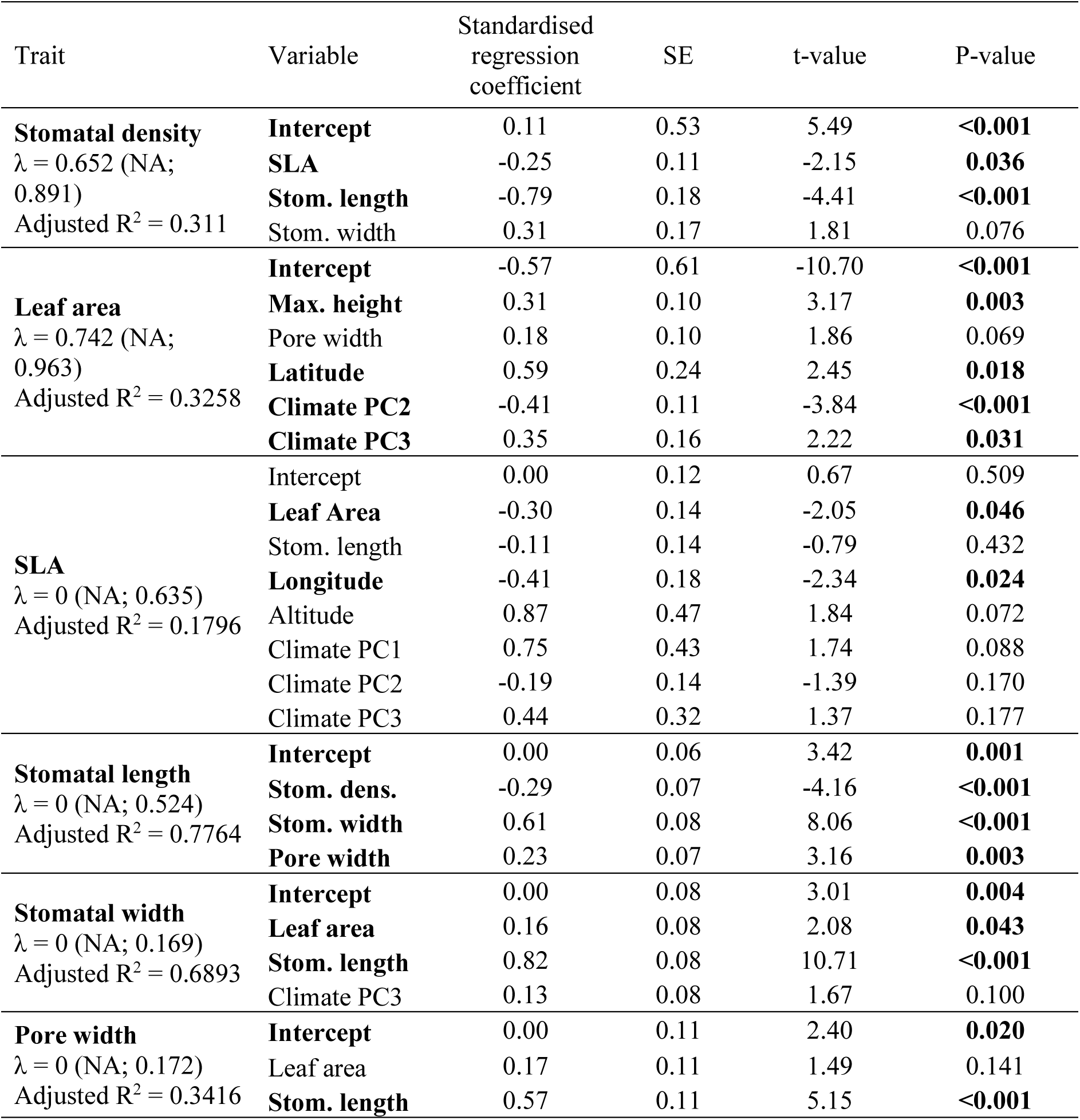
Multiple PGLS regressions fitted by maximum likelihood across *Coffea* without outgroups for each trait. Significant predictors are marked in **bold**. For stomatal density, leaf area, and SLA, n = 58; for stomatal length, stomata width, and pore width, n = 55.

### Relationships among leaf traits

Across all examined *Coffea* species, PGLS regressions showed that stomatal density was significantly negatively related to SLA (t_51_ = -2.15, P = 0.036) and stomatal length (t_51_ = -4.41, P < 0.001), with a marginally significant positive relation to stomatal width (t_51_ = 1.81, P = 0.076). Leaf area was significantly positively related to maximum plant height (t_49_ = 3.17, P = 0.003), and a marginally significant positive effect of pore width on leaf area was also detected (t_49_ = 1.86, P = 0.069). For SLA, the best regression model revealed a negative relationship with leaf area (t_47_ = -2.05, P = 0.046). Stomatal length was negatively predicted by stomatal density (t_51_ = -4.16, P < 0.001). Also, stomatal width (t_51_ = 8.06, P < 0.001) and pore width (t_51_ = 3.16, P = 0.003) both had significantly positive effects on stomatal length. Leaf area (t_51_ = 2.08, P = 0.043) and stomatal length (t_51_ = 10.71, P < 0.001) were the only significant predictors of stomatal width, both having a positive effect. Finally, pore width was significantly related to only one independent variable: stomatal length (t_52_ = 5.15, P < 0.001), of which the effect was positive.

Regressions restricted to the EC clade showed qualitatively similar results to the regressions across the genus for trait-trait relationships. Significant predictors differed only for SLA, where a marginally significant negative relationship with maximum plant height (t_46_ = - 1.86, P = 0.069) was observed. Differences became more apparent when further narrowing the analyses to the AFR and WIOI subclades. Leaf area was positively related to stomatal density in the AFR subclade (t_14_ = 3.51, P = 0.003), which was not the case in any other clade. Leaf area was also positively related to maximum plant height in the WIOI subclade (t_28_ = 2.59, P = 0.015). Further, the relationship between SLA and leaf area - which was positive across the genus - was significantly negative across the AFR subclade (t_15_ = 2.60, P = 0.020), and SLA was inversely related to maximum plant height (t_15_ = -5.44, P < 0.001) in the AFR subclade. Finally, stomatal width was significantly positively related to maximum plant height in the AFR subclade (t_15_ = 3.83, P = 0.002).

### Relationships between leaf traits and environment

From the phylogenetic PCA applied to the 19 bioclimatic variables, three PCs were retained and added to the data set. The three retained PCs jointly explained 79% of the variance in bioclimatic variables. PC1 and PC2 explained 35% and 31% of the variance, respectively (Figure 3, **[Supplementary information]** Table S5). PC1 represented an axis of temperature, with higher values indicating hotter climates. The strongest loadings for PC2 were all negatively related to precipitation. This axis thus generally represented an axis of drought, with higher PC2 values indicating less rainfall. The third PC was interpreted as an axis of temperature seasonality, where higher PC3 values indicate stronger seasonal temperature fluctuations.

**Figure 3:**
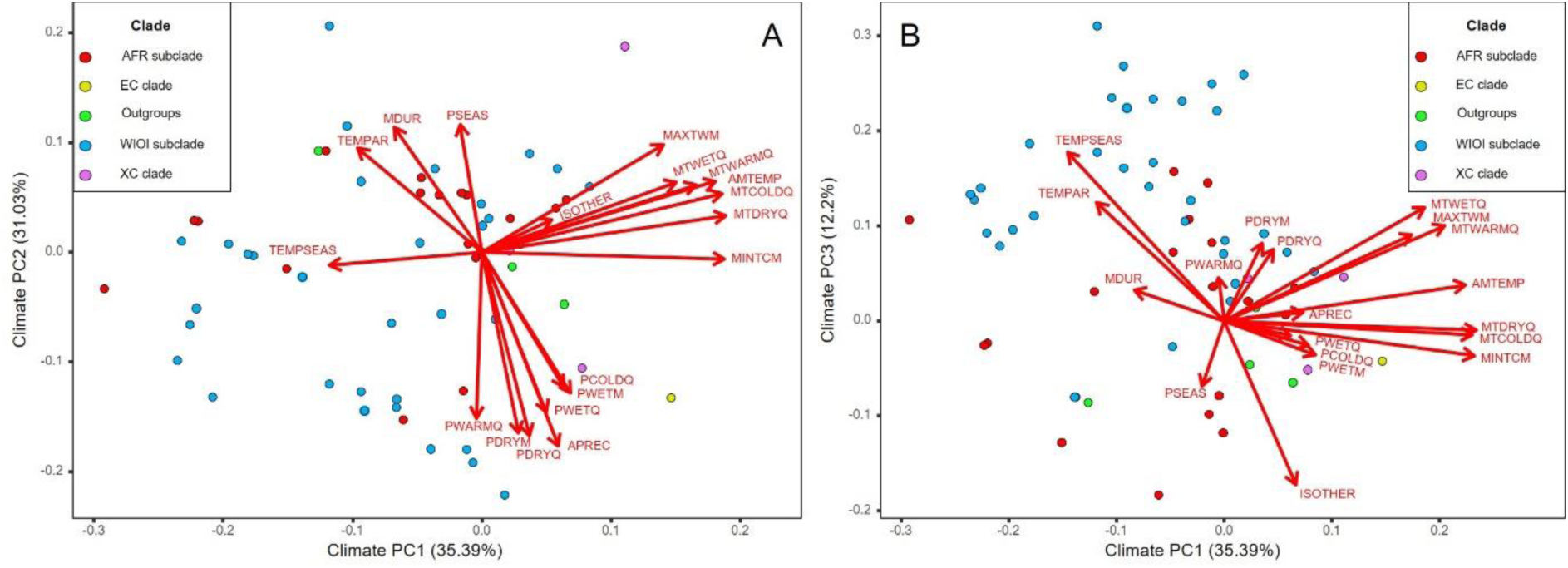
Principal component analysis of the 19 bioclimatic variables drawn from WorldClim (Fick & Hijmans, 2017). Climate variable abbreviations are explained in **[Supplementary information] Table S5**. Data points indicate species (n = 62) and are color-coded according to their clade. **A**, PC1 versus PC2; **B**, PC1 versus PC3. AFR = African; EC = Eu-coffee; WIOI = West Indian Ocean Islands; XC = Xeno-coffee. Clade nomenclature is based on Hamon et al. (2017).

Significant relationships between environmental variables and leaf traits were uncommon across the genus (Table 2). For stomatal density and the three stomatal dimensions, no significant relations with environmental variables were detected (though climate PC3 was retained as a non-significant predictor in the best-fitting model for stomatal width). For SLA, all three climate PCs as well as altitude and longitude were retained in the final model, but only the negative relation to longitude was significant (t_47_ = -2.34, P = 0.024). The positive effects of altitude (t_47_ = 1.84, P = 0.072) and climate PC1 (t_47_ = 1.74, P = 0.088) on SLA were marginally significant. The only leaf trait strongly affected by our environmental variables was leaf area, with significant positive relations to latitude (t_49_ = 2.45, P = 0.018) and climate PC3 (t_49_ = 2.22, P = 0.031), and a strong negative relation to climate PC2 (t_49_ = -3.84, P < 0.001).

Trait-environment associations across the EC clade were similar, with only predictors of SLA showing qualitative differences. The association between longitude and SLA was no longer significant in the EC clade, while the positive relation between SLA and altitude (t_46_ =2.31, P = 0.026) and climate PC1 (t_46_ = 2.28, P = 0.027) became significant. As was the case for trait-trait relationships, patterns differed substantially when looking only at the AFR or WIOI subclades. Whereas stomatal density was not related to climate at a larger phylogenetic scale, it showed significant negative associations with altitude (t_11_ = -7.16, P < 0.001), climate PC1 (t_11_ = -7.87, P < 0.001), and climate PC3 (t_11_ = -2.81, P = 0.017) in the AFR subclade, and positive associations with altitude (t_26_ = 2.62, P = 0.014) and climate PC3 (t_26_ = 4.75, P < 0.001) in the WIOI subclade. Leaf area showed significant positive relations with altitude (t_14_ = 3.12, P = 0.008) and climate PC1 (t_14_ = 4.02, P = 0.001) in the AFR subclade and a negative association with climate PC2 (t_28_ = -2.63, P = 0.014) across the WIOI subclade. SLA was not related to climate or geography in the AFR subclade. In contrast, analysis of SLA in the WIOI subclade uncovered a significant negative relationship with latitude (t_28_ = -3.15, P = 0.004) and positive associations with altitude (t_28_ = 3.17, P = 0.004) and climate PC1 (t_28_ = 2.85, P = 0.008). For stomatal length, significant environmental associations were only detected in the AFR subclade, all of which were negative: longitude (t_11_ = -3.97, P = 0.002), climate PC2 (t_11_ = -2.67, P = 0.022), and climate PC3 (t_11_ = 3.08, P = 0.011). Stomatal width and pore width showed no environmental association in any of the tested clades.

### Evolutionary model comparisons

Model comparisons revealed that OU models outperformed the BM model for all traits, but the WN model was a significantly better fit for stomatal density, stomatal width, and pore width (ΔAICc > 2, Table 3, **[Supplementary information]** Table S6). Leaf area was best explained by the OU_D_ model (AICc = 139.8), significantly outperforming all other models (ΔAICc > 2). For SLA, the OU_S_ model (AICc = 317.2) was the best fit, slightly outperforming the OU_1_ model (AICc = 318.6). Stomatal length showed the lowest AICc value for the OU_1_ model (AICc = 281.2). Finally, PGLS regression models displayed significantly lower AICc values than the evolutionary models for all traits except SLA (Table 3).

**Table 3:**
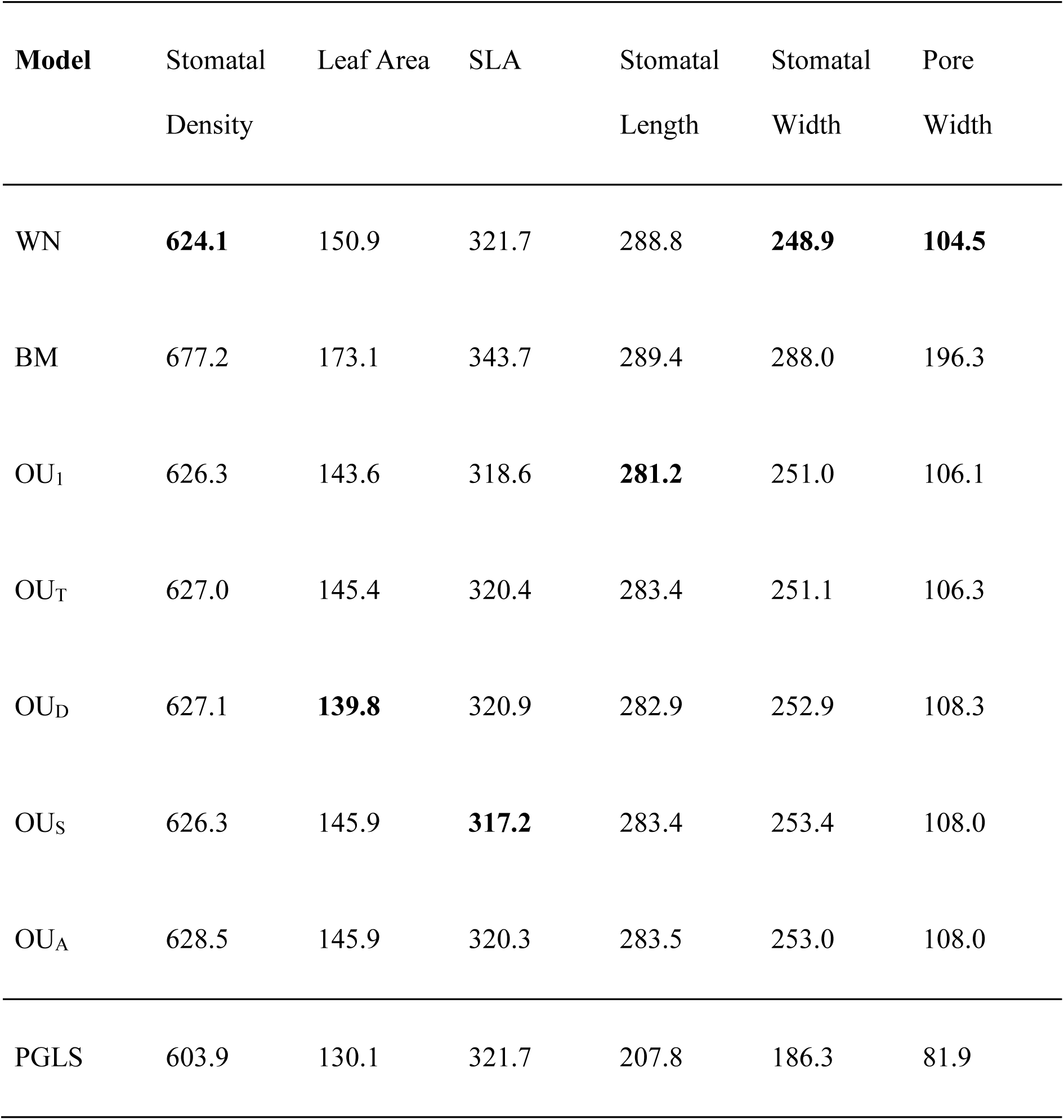
Model AICc values across *Coffea* without outgroups for each trait. The lowest AICc value for each trait (not considering the PGLS models) is marked in **bold**. For stomatal density, leaf area, and SLA, n = 58; for stomatal length, stomata width, and pore width, n = 55.

## Discussion

Leaf functional traits play a key role in environmental adaptability. Their variation and evolution across a genus can shed light on the factors driving evolutionary change, and on the adaptive value of specific traits. This information can be of use in strategies for conservation and crop improvement. Our results confirm that *Coffea* species have diverged substantially in the measured leaf functional traits over evolutionary time, contributing to the mapping of trait variation in the genus. Intraspecific trait differences were less substantial compared to interspecific differences for five of the six measured traits, and traits evolved in a correlated fashion at different taxonomic levels. It was also clear that climate played an important role in driving the evolution of some leaf functional traits in *Coffea* and that phylogenetic patterns in these traits differed between subclades.

### Genus-wide leaf trait patterns

Our results suggest that leaf traits in *Coffea* have evolved in a rather continuous mode, from low densities of large stomata in the outgroup and early branching XC clade to higher densities of smaller stomata in the EC clade. This clear trade-off between stomatal size and density has been shown in other taxa (e.g., Doheny-Adams et al., 2012; Brodribb et al., 2013), and was evident in our trait PCA and PGLS regressions. More early branching taxa, such as the XC clade and *C. charrieriana,* tend to have fewer, larger stomata than more recently derived species such as e.g., *C*. *mauritiana*. This observation may suggest a shift in trait evolution throughout the diversification of the genus. However, we detected no clear climatic drivers of this shift, indicating that it may have been driven by variables not considered in this study. As higher densities of smaller stomata have been associated with increased stomatal conductance (Franks & Beerling, 2009; Drake et al., 2013), we hypothesise that this shift may have been a response to steadily decreasing atmospheric [CO_2_] throughout the diversification of *Coffea* over the past 12 million years (Hamon et al., 2017; Rae et al., 2021; Haworth et al., 2023). Lower [CO_2_] could have led species to increase their stomatal conductance to maintain the same growth rates, explaining the trend observed here.

Unlike stomatal size and density, SLA and leaf area were poorly related, both across the genus and within the subclades. Contrasting relationships across clades and the relatively unrelated environmental associations of these traits support the conclusions of Ackerly et al. (2002) that these traits are related to different aspects of plant performance in an environmental context. Further, leaf area increases with maximum plant height across the genus *Coffea*, an observation that aligned with our expectations based on literature (Jensen & Zwieniecki, 2013; Price et al., 2014; Peel et al., 2017). The work of Price et al. (2014) showed that traits in the LES do not tend to vary with maximum plant height across species, except for leaf area.

When considering all included *Coffea* species or the EC clade, leaf area was strongly negatively related to drought, and positively to latitude. Species with larger leaves thus occur in environments with more rainfall and at more northern latitudes, i.e., in more equatorial regions, as has been observed in other taxa (e.g., Givnish, 1984; Peppe et al., 2011). We hypothesised that higher aridity would select for smaller leaf sizes to limit water loss via transpiration (Poole & Miller, 1981; Thuiller et al., 2004; Wright et al., 2017; Chaturvedi & Raghubanshi, 2018). For example, in a study on the genus *Leucadendron* in the Cape Floristic region, Thuiller et al. (2004) observed significantly smaller leaves in species in arid environments than in species in more moist habitats. The observed patterns thus aligned with our initial expectations, reflecting a functional trade-off between drought tolerance and photosynthetic productivity (Thuiller et al., 2004).Overall, trait-climate relationships across the genus were relatively weak, as we had hypothesised. It is likely that trait evolution is influenced by multiple factors. Environmental factors also differ strongly across the broad geographic range of the genus, making it difficult to discern direct drivers across *Coffea* as a whole.

### Contrasting patterns between subclades

Both for trait-trait and trait-environment relationships, we observed varying patterns depending on the clade considered. These differences highlight the importance of clade-specific analyses, especially when clades are also geographically isolated.

Stomatal density exhibited strong positive relationships with leaf area and with maximum plant height in the AFR subclade species studied here, relationships that were not present (or not as prominent) in the WIOI subclade. Taller species and species with larger leaves thus tend to have more stomata per mm^2^ in the AFR subclade, a result that is contrary to observations in other taxa (Kouwenberg et al., 2007; Carins Murphy et al., 2012; Peel et al., 2017; Conesa et al., 2020). For example, Carins Murphy et al. (2012) studied the subtropical tree species *Toona ciliata* and observed lower stomatal density in larger leaves, but they additionally considered irradiance levels as a defining factor. According to their findings, more shaded plants (i.e., shorter species) have larger leaves due to the expansion of epidermal cells, leading to lower stomatal densities. This supports our observation that shorter species have lower stomatal densities, but the positive association between leaf size and plant height (in contrast with Carins Murphy et al., 2012) excludes the possibility of a leaf size-mediated relationship between plant height and stomatal density. The cause of our contrarian findings requires further study.

The negative relationships between stomatal density and both temperature and altitude in the AFR subclade indicate that species in cooler environments have higher stomatal densities. This observation is again in direct contrast to patterns detected in other temperate (Kouwenberg et al., 2007) and subtropical (Liu et al., 2020) species, though some studies have found ambiguous or no patterns in this relationship in both (sub)tropical (Hill et al., 2014; Zhao et al., 2016) and temperate (Wang et al., 2014) climates. For example, Liu et al. (2020) observed increasing stomatal densities with elevation in three subtropical mountain species, whereas Zhao et al. (2016) observed no relationship between stomatal density and altitude across 105 angiosperm species in (sub)tropical montane forests in Yunnan (South-west China). The relation between stomatal density and the environment may thus differ across taxa and across habitats. On the African continent, species in warmer environments and/or at lower altitudes show a higher density of smaller stomata, thereby accelerating the rate of gas exchange and providing more potential for evaporative cooling (Drake et al., 2013; Hill et al., 2014). The same functional explanations could apply to the WIOI subclade, where the positive relationship between stomatal density and temperature seasonality aligns with the observations of Drake et al. (2013): higher stomatal densities (and thus smaller stomata) have more dynamic stomatal characteristics and may therefore be better adapted to respond to temperature fluctuations throughout the year (Drake et al., 2013).

SLA was also negatively related to maximum plant height in the AFR subclade. A core trait of the LES, high SLA values indicate proportionally lower levels of leaf investment (Wright et al., 2004; Reich, 2014). Taller species are generally more exposed to higher levels of solar irradiance, whereas shorter plants tend to be more shaded. Shaded leaves are known to have higher SLA values than sun leaves to enhance light capture in the shade (e.g., Niinemets, 2010; Legner et al., 2014; Paź-Dyderska et al., 2020), a pattern which appears to hold across the AFR subclade of *Coffea*. The fact that we did not observe this relationship in the WIOI subclade is likely due to the more homogeneous distribution of plant height in this subclade (results not shown) or due to the wider variety of vegetation structures inhabited by Madagascan species (Davis et al., 2006).

Leaf area was positively related to altitude and temperature in the AFR subclade, but not in the WIOI subclade. Leaf size has often been related to temperature (Ackerly et al., 2002; Royer et al., 2005; Wright et al., 2017; Glade-Vargas et al., 2018), though patterns are not consistent across taxa or taxonomic levels and depend on other factors such as rainfall or solar irradiance. Larger leaves have a higher potential for evaporative cooling, though this cooling requires sufficient water availability and should thus be conditional on sufficiently high moisture levels.

### Evolutionary processes shaping leaf trait evolution

The non-uniform distribution of leaf trait values across the genus emphasises the importance of a phylogenetic perspective when evaluating trait differences. Our results show that a pure drift BM model of evolution poorly predicted leaf functional trait evolution across *Coffea*.

Though univariate estimates showed significant phylogenetic signal for SLA and stomatal length across *Coffea*, multivariate phylogenetic signal was not significantly different from zero for any trait across the genus. An intrinsic property of OU models (and thus also of WN models; Münkemüller et al., 2015) is that evolutionary history becomes gradually less important over time, reducing phylogenetic signal (Felsenstein, 1988; Blomberg et al., 2003; Ackerly, 2009). However, our results showed that modes of trait evolution differed among subclades. In the AFR subclade, multivariate phylogenetic signal λ was high and was not significantly different from 1 for stomatal density and stomatal length. In the WIOI subclade, λ was also high for stomatal density, but stomatal length showed no phylogenetic signal. These differences may be partially explained by the different evolutionary trajectory of these clades, such as a rapid radiation of the WIOI clade across Madagascar & the Mascarenes (Anthony et al., 2010).

The evolution of stomatal density, stomatal width, and pore width was better explained by a phylogeny-independent WN model than by any of the phylogenetic models fitted in this study. This could be due to rapid evolution along the phylogeny, or due to strong environmental influences on these traits (Glade-Vargas et al., 2018; Capunitan et al., 2020). The latter may be the case for stomatal density, which showed strong but differing associations with climate in each subclade.

The evolution of stomatal length was best approximated by the OU_1_ model, suggesting stabilising selection towards a single optimum value for the entire genus. The best-fitting model for leaf area was the OU_D_ model, whereas the OU_S_ model provided the best fit for SLA. Evolutionary divergence in these three traits was thus constrained compared to a pure drift BM model, which is supported by the absence of multivariate phylogenetic signal and the high observed rates of adaptation. Moreover, leaf area and SLA appeared to be driven by variation in climate. In the case of leaf area, this interpretation is supported by the PGLS regressions: leaf area was negatively related to climate PC2 across all clades except the AFR subclade. We can thus state with confidence that the evolution of leaf area across *Coffea* is driven by variation in precipitation experienced by the different species. Conversely for SLA, relationships observed in the PGLS regressions do not appear to corroborate the trait’s evolution in response to temperature seasonality. Climate PC3 was not a significant predictor of SLA in any of the tested clades, though it was retained in the best-fitting models across *Coffea* and across the EC clade. Moreover, the difference in fit between the OU_S_ and OU_1_ models for SLA was minimal and nonsignificant (ΔAICc = 1.4). We are therefore more hesitant to state that the evolution of SLA has been driven by temperature seasonality in *Coffea*, but we did observe constrained evolution and thus stabilising selection on SLA within the genus.

For all traits except SLA, the AICc value of the PGLS model was substantially lower than any of the other models, indicating that the inclusion of trait-trait and trait-environment associations explained a meaningful proportion of trait variation that was not considered in the evolutionary WN, BM, and OU models. The benefit of these evolutionary models over PGLS is that they are explicitly evolutionarily interpretable, whereas PGLS models are simply regression models correcting for trait similarity resulting from common ancestry (Hansen, 1997).

Our results suggest that trait diversification across *Coffea* is substantially constrained compared to the expectation under pure drift (BM) evolution, based on evolutionary model comparisons and multivariate estimates of phylogenetic signal. This ecological similarity among related species can result from multiple processes, however, including insufficient variation, gene flow, genetic constraints, or stabilising selection (Wiens & Graham, 2005; Losos, 2008; Mitchell et al., 2018). Our analyses show that two highly derived sister species exhibit markedly different leaf traits: *C. mauritiana* and *C. macrocarpa* (Figure 1B). The former was sampled on the island of Réunion, whereas the latter is endemic to Mauritius (Nowak et al., 2014). Their strong trait divergence despite their phylogenetic proximity suggests that isolation and novel niche availability following dispersal led to rapid trait evolution and divergence. Thus, rapid evolution is possible within the genus, and we deem strong genetic constraints unlikely to inhibit trait divergence across *Coffea*. Gene flow may occur at small scales between species but is unlikely to constrain evolutionary divergence at the genus level, or at the wide spatial scale considered here. We therefore consider stabilising selection to be the driving force behind the constrained evolution of the considered leaf traits across the genus. Leaf traits are known to be more labile and adapt rapidly relative to other plant traits, such as seed or floral traits, which are generally more evolutionarily conserved (Ackerly, 2009; Vandelook et al., 2018). Strong stabilising selection would thus act efficiently on the considered leaf traits, exemplified by the absence of phylogenetic signal and the high rates of adaptation. In contrast to our observation that leaf traits are evolutionarily labile, Glade-Vargas et al. (2018) detected significant phylogenetic signal for 14 out of 20 leaf traits in the Nothofagaceae family, indicating that these leaf traits were evolutionarily conserved. However, they considered only univariate estimates of phylogenetic signal and did not account for environmental effects.

### Methodological limitations

Because of statistical limitations in fitting OU models on small phylogenies (Beaulieu et al., 2012; Cooper et al., 2016), we did not consider evolutionary model comparisons in the AFR and WIOI subclades. We also chose not to consider more complex models of continuous trait evolution (e.g., mvSLOUCH; Bartoszek et al., 2012) for the sake of interpretability. These models may nonetheless provide useful insights into the mechanisms and drivers of *Coffea* leaf trait evolution and should be considered in future studies.

Due to our exclusive use of herbarium specimens, the trait variation detected in this study is likely an underestimation of the variation present in wild populations and may suffer from unstandardised sampling, particularly in WIOI species which occur in a broad range of vegetation types (Davis et al., 2006). Contrarily, since most central African *Coffea* species grow in the rainforest understory, phenotypic variation due to variation in light exposure can be considered minimal in central African species (Hatangi et al., 2023). Despite these limitations, the use of herbarium accessions allows for studies at a large geographical scale including almost half of all known *Coffea* species, which would not be feasible otherwise. Nonetheless, more sampling and improved data on habitat, vegetation structure, and local climate remains necessary.

## Conclusion

Our study seems to indicate that the evolution of leaf functional traits has been a complex process in *Coffea*. Interrelations with other traits and with various environmental variables, differing across clades, make it difficult to disentangle individual drivers of trait evolution. We show that the genus *Coffea* exhibits substantial leaf functional trait variation, and we observed a directional shift from low densities of large stomata towards higher densities of smaller stomata, potentially driven by a decline in atmospheric [CO_2_] during the divergence of *Coffea*. Also, by incorporating both ecological variables and explicit evolutionary models in our analyses, we found that the evolution of leaf area was most likely driven by differences in precipitation across *Coffea*. We could confirm that leaf traits tend to adapt rapidly, with an important role for stabilising selection, and that trait evolution across *Coffea* was not governed by pure drift. Contrasting correlations and differing evolutionary processes across different clades also highlight the relevance of a phylogenetic perspective in comparative studies, showing that the strength and direction of trait relationships depends on the considered taxonomic scope and geographic range. Our study can fundamentally be of relevance to crop improvement programmes, as our results suggest that smaller-leaved species are adapted to drier climates. Therefore, breeding programmes may consider leveraging leaf area in climate adaptation strategies. Our findings contribute more broadly to our understanding of the variation and adaptive significance of leaf functional traits in *Coffea*.

## Supplementary Information

Supplementary Information can be found online and consists of the following files:

Table S1: individual leaf data set

Table S2: Species-level data set

Table S3: loadings of the trait PCA

Table S4: Regression tables and other output of PGLS across all considered clades

Table S5: loadings of the climate PCA

Table S6: parameter output of the evolutionary models Figure S1 (A-F): Boxplots of trait values across the genus

## Funding

This work was financially supported by an FWO research project grant (G090719N), and by the Belspo funded COBECORE project (R/175/A3/COBECORE).

## Acknowledgements

We would like to thank Mieke Rotteveel, Sofie Meeus, and Martine Borremans for help with data collection and Ann Bogaerts for facilitating access to the BR herbarium.

## Conflict of interest

We report no conflicts of interest.

## Appendix Shrinkage correction

To correct for potential bias introduced in SLA values by leaf shrinkage of herbarium specimens, 145 fresh leaves were sampled without petioles across 29 live accessions of 11 *Coffea* species. These leaves were weighed using a precision scale (1 mg accuracy) and scanned with a flatbed scanner (Canon CanoScan 9000F Mark II) before and after fully drying in a herbarium oven. Fresh and dry leaf area were measured with ImageJ open access software version 1.53v (Schneider et al., 2012), after which leaf shrinkage factors were calculated as the ratio of dry area to fresh area. We then used dry area, which has been found to be a good predictor of shrinkage (Blonder et al., 2012), to predict shrinkage factors for all other leaves in the data set using a simple linear regression: (*Shrinkage factor = 0.008851 ∗ log_e_(Dry area) + 0.895520*, P < 0.001, though residuals deviated from normality: Shapiro-Wilk normality test, W = 0.87496, P < 0.001). Data on SLA (calculated using dry area) was divided by the obtained shrinkage factors, resulting in SLA data with a correction for leaf shrinkage.

## Notes

### Competing Interest Statement

The authors have declared no competing interest.

