## Supplemental Figure S1 for "Leaf functional trait evolution and its putative climatic drivers in African *Coffea* species"

### Slide 1
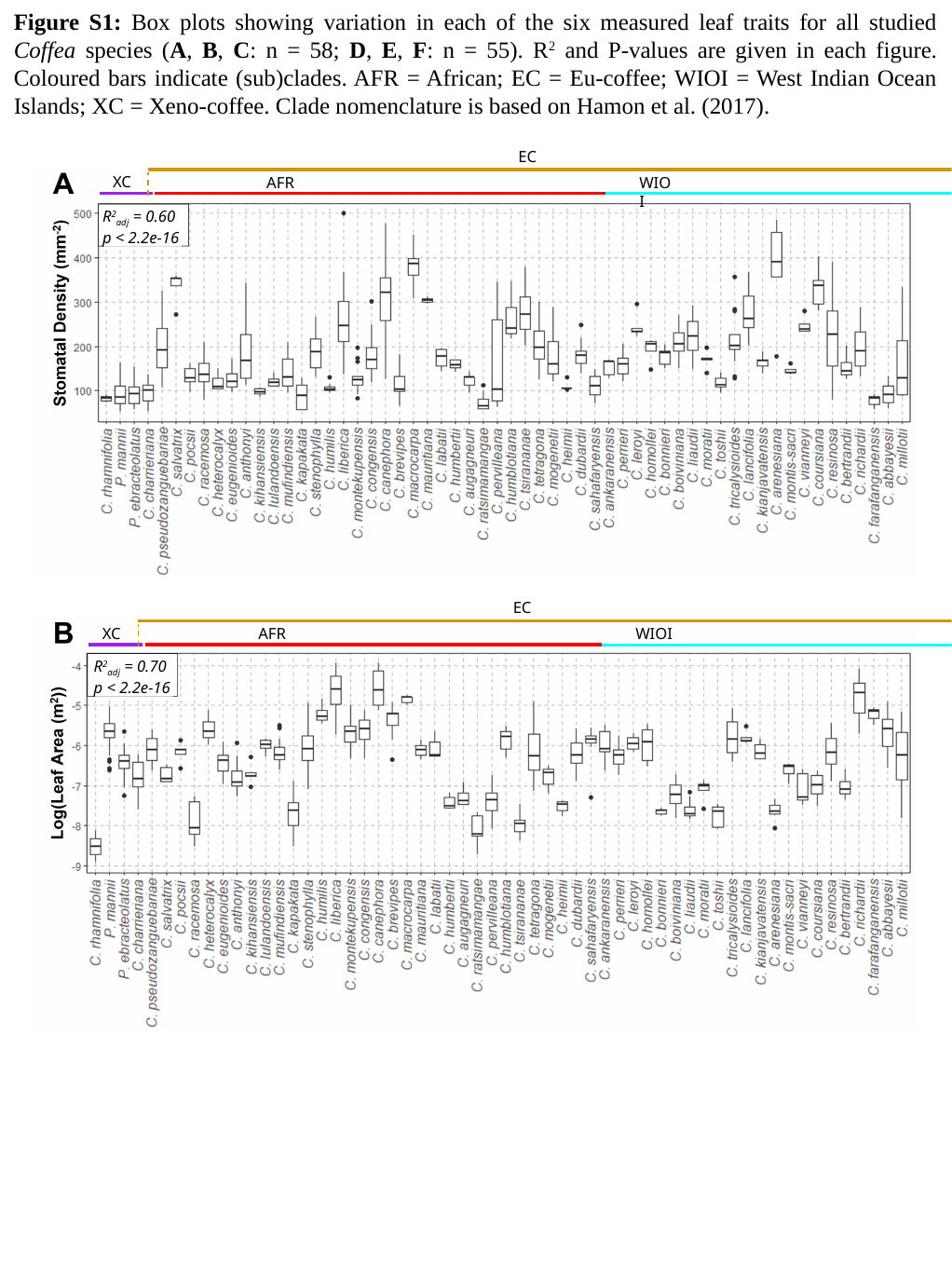

Figure S1: Box plots showing variation in each of the six measured leaf traits for all studied Coffea species (A, B, C: n = 58; D, E, F: n = 55). R^2^ and P-values are given in each figure. Coloured bars indicate (sub)clades. AFR = African; EC = Eu-coffee; WIOI = West Indian Ocean Islands; XC = Xeno-coffee. Clade nomenclature is based on Hamon et al. (2017).
EC
XC
AFR
WIOI
R^2^_adj_ = 0.60
p < 2.2e-16
EC
XC
AFR
WIOI
R^2^_adj_ = 0.70
p < 2.2e-16

### Slide 2
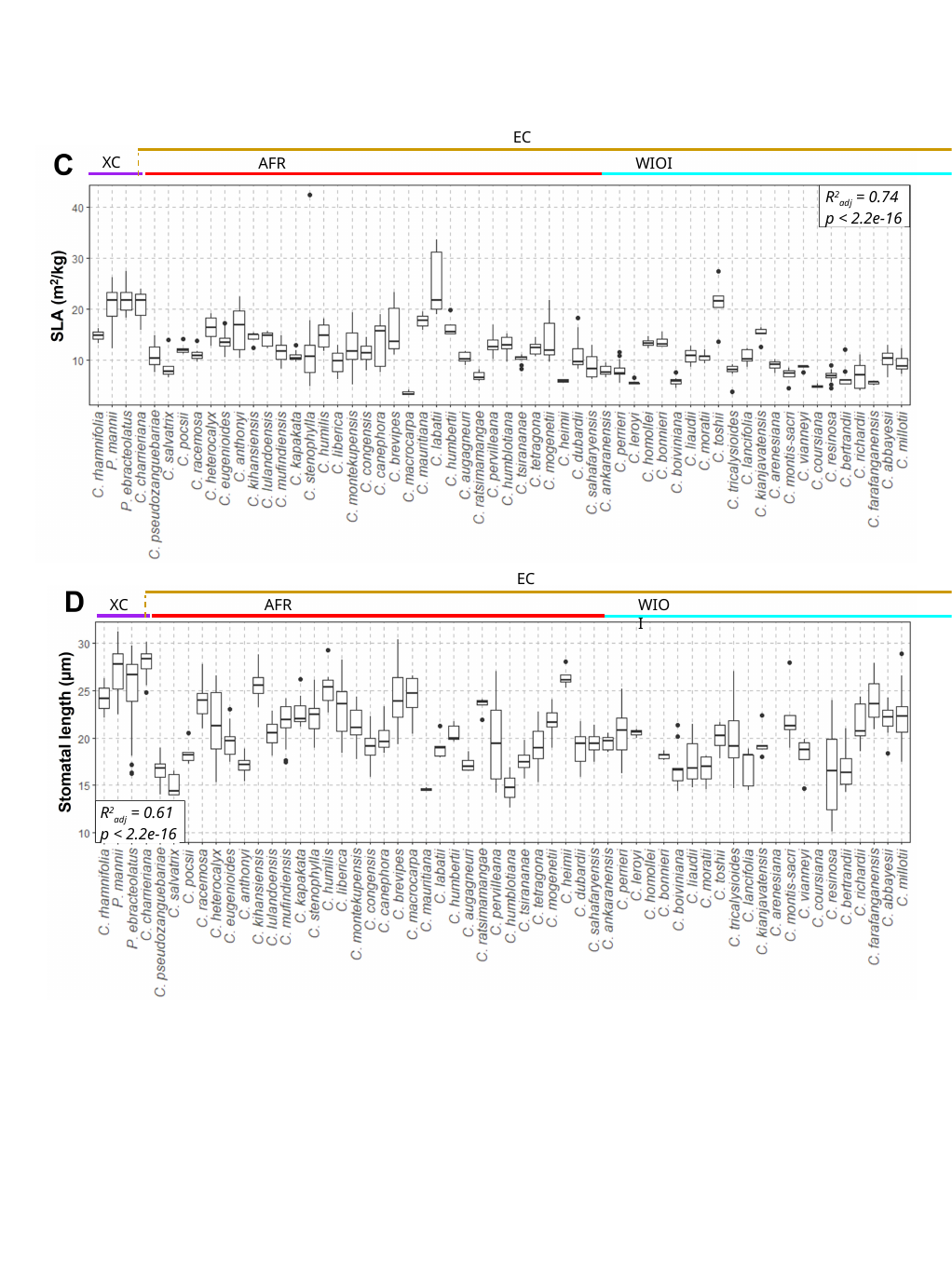

EC
XC
AFR
WIOI
R^2^_adj_ = 0.74
p < 2.2e-16
EC
XC
AFR
WIOI
R^2^_adj_ = 0.61
p < 2.2e-16

### Slide 3
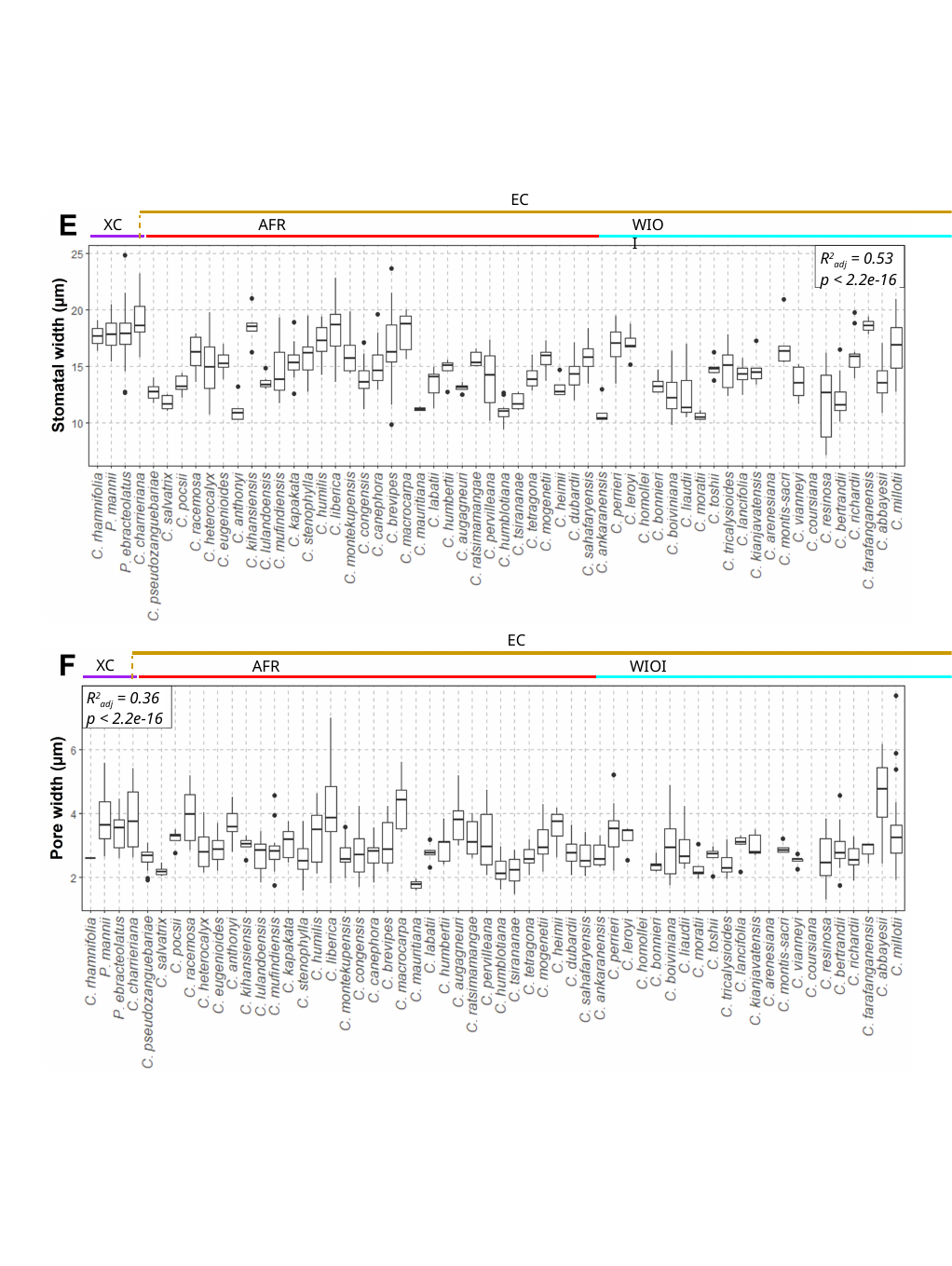

EC
XC
AFR
WIOI
R^2^_adj_ = 0.53
p < 2.2e-16
EC
XC
AFR
WIOI
R^2^_adj_ = 0.36
p < 2.2e-16
